# Embedded Printing-Derived Pyrolytic Carbon Electrodes with Graphene-like Features for Neural Stem Cell Differentiation and Dopamine Sensing

**DOI:** 10.64898/2026.10.04.756553

**Authors:** Hakan Gürbüz, Asli Aybike Dogan, Babak Rezaei, Giorgio Scordo, Thomas Willum Hansen, Janko Kajtez, Marta P. Pereira, Stephan Sylvest Keller, Martin Dufva, Arto Heiskanen, Jenny Emneus

## Abstract

Carbon is a versatile electrode material that is used in applications ranging from electrical energy storage and electrosynthesis to CO_2_ conversion, H_2_ production, and biosensing. The development of pyrolytic carbon electrodes derived from lithographically patterned photoresist structures has expanded the possibilities for creating diverse 2D and 3D electrode designs. In this paper, we present disc-shaped SU-8-derived pyrolytic carbon scaffold electrodes with an open hexagonal honeycomb lattice. The scaffolds were fabricated, using embedded 3D printing within a shear-thinning support bath composed of fused silica nanoparticles (7% w/v) in mineral oil. The rheological properties of the support bath were optimized to enable the printing of low-viscosity materials like SU-8, which would otherwise lack structural integrity in standard extrusion-based 3D printing. The SU-8 scaffolds were then pyrolyzed and characterized using SEM, Raman spectroscopy, Energy Dispersive X-ray, X-ray Photoelectron Spectrometry, and Transmission Electron Microscopy, which revealed graphene-like ‘microflowers’ on its surface. Electrochemical analysis demonstrated excellent electrochemical properties. The functionality of the scaffold electrodes was validated by differentiation of human neural stem cells into dopaminergic neurons on the scaffold surface, followed by real-time detection of dopamine exocytosis upon cell depolarization.

Supplementary information for this article is available on-line.

## 1. Introduction

Carbon, in its various forms, is a widely used electrode material for electrochemical detection owing to its excellent electrochemical surface activity, broad potential window, chemical stability, and high capacity for surface modification and functionalization [1]. Traditional methods, such as screen-printing and inkjet printing, offer simple, low-cost, and flexible ways to produce 2D carbon electrodes [2–5]. Carbon microelectromechanical systems (CMEMS) emerged in the 1970s and rely on photolithographic patterning of polymer precursor templates followed by high-temperature pyrolysis (above 900°C in an inert atmosphere) to create precise, reproducible 2D and 3D carbon microelectrodes [6,7]. While CMEMS is more expensive than inkjet or screen-printing, it allows for intricate 3D structuring of carbon electrodes with various geometries such as pores, cavities, pillar arrays, and micro/nanograss, all of which enhance sensitivity by increasing the surface area available for electrochemical detection [8–10]. However, CMEMS is limited to small structures and photocurable resins.

3D printing emerges as a promising alternative for scaling up the fabrication of pyrolytic carbon structures from any pyrolyzable polymer precursor [11]. Here, we present a method that uses embedded printing of SU-8 in a sheer-thinning liquid (support bath), composed of a mixture of mineral oil and fumed silica nanoparticles, which supports and stabilizes the printed 3D SU-8 structure. This technique enables omni-directional printing of non-viscous liquids, such as SU-8, which otherwise cannot support their own weight [12,13]. The printed structures are subsequently cured and removed from the support bath, followed by pyrolysis into 3D carbon electrodes

We previously used fused deposition modelling (FDM) to fabricate disc-shaped polycaprolactone (PCL) scaffolds with an open hexagonal honeycomb lattice for cerebral organoid engineering, resulting in organoids exhibiting gyrification-like folding patterns [14]. Building on this design, the aim of the present study was to develop disc-shaped SU-8-derived pyrolytic carbon scaffold electrodes with dual functionality. These scaffolds were designed to provide structural support for neural cell growth and differentiation while simultaneously enabling electrochemical monitoring of neurotransmitter release. As a proof of concept, human neural stem cells (hNSCs) were differentiated into dopaminergic neurons, and exocytotic dopamine release was detected electrochemically [15]. This work establishes the foundation for future electrically active scaffold platforms for brain organoid applications.

The same basic computer-aided design (CAD) model as reported in[14] was used to fabricate disc-shaped SU-8 scaffolds with an open hexagonal honeycomb lattice via embedded 3D printing. Subsequent pyrolysis converted the polymeric scaffolds into conductive pyrolytic carbon scaffold electrodes while preserving the overall architecture. The resulting structures were characterized for their morphology, chemical composition, and electrochemical properties, including the formation of graphene-like carbon microflowers on the scaffold surface during pyrolysis. Finally, the functionality of the scaffold electrodes was demonstrated by differentiation of human neural stem cells (hNSCs) into a dopaminergic phenotype and enabling electrochemical detection of dopamine released following chemically induced exocytosis.

## 2. Method

### 2.1 Preparation and characterization of fumed silica nanoparticle-mineral oil support bath

Preparation of the support bath was adapted with slight changes from the approach presented by Jin et al.[12] Commercial hydrophobic fumed silica nanoparticles with an average diameter of 150 nm (Aerosil@R 972 Pharma from Evonik Industries AG, Germany) were used. Different support baths were prepared bydispersing the appropriate amount of dry fumed silica powder in light mineral oil (Sigma-Aldrich) at room temperature to obtain suspensions of 6, 7, and 8 % (w/v). After continuous mixing overnight using a magnetic stirrer at 400 rpm, the support baths were aged overnight and then centrifuged at 3000 rpm for 15 minutes to thoroughly eliminate any trapped bubbles. Rheological characterization was performed based on adaptation of the protocol presented by Chen et al [16] using a Discovery Hybrid Rheometer 2 (TA instruments, New Castle, Delaware) equipped with a 40 mm parallel steel plate and a steel Peltier plate. Dynamic strain sweeps were conducted in the range of 0.01 to 500 %. Dynamic frequency sweeps were performed at a low strain of 0.2 %. Shear rate sweeps were performed by varying the shear rate from 0 to 50 Hz. During transient step shear rate tests, the support baths at different concentrations were sheared at 10 Hz and a low strain of 0.2 % for 120 s and at an increased strain of 500 % for 60 s. The viscosity variation was shown during five cycles of the same step order. Temperature sweep was conducted by raising the temperature from 25 to 100 °C at a rate of 1°C/10 s and the storage modulus measured. The UV transparency of the supports baths was determined spectrophotometrically (Spark Multimode Microplate Reader, Tecan, Sweden) by scanning the wavelength from 200 to 1000 nm.

### 2.2 Design and fabrication of SU-8 derived carbon scaffold electrodes

Scaffolds with honeycomb patterns were designed using SolidWorks 2020 (Dassault Systems) 3D modelling software and exported as stereolithography (STL) files. The STL files were imported into Simplify3D slicer software (Simplify3D Inc.) to create the print path. An extrusion-based 3D bioprinter (Felix BIO, FELIXprinters B.V.) was used as the dispensing system. A 30-gauge metal blunt-end dispensing tip was used for SU-8 printing (Fisnar, USA). For embedded printing of SU-8 (negative photoresist SU-8-2035, Microchem Corp.), the 7 % (w/v) fumed silica nanoparticle-mineral oil support bath was used. The 3D printed SU-8 scaffolds were UV cured while still inside the support bath for 15 min followed by heat treatment in a convection oven at 60 °C for 1 h. The cured scaffolds were removed from the gel and washed in 2-propanol and toluene (both from VWR International, LLC, part of Avantor, Inc, USA) to remove the leftover gel. The cleaned scaffolds were then developed in a SU-8 developer solution (Propylene glycol methyl ether acetate, Sigma Aldrich). Finally, the developed scaffolds were pyrolyzed in a PEO-604 furnace (ATV Technology, Germany) under N_2_ gas flow (200 sccm) at 900 °C. The temperature was ramped up at a rate of 2 °C/min until 410 °C was reached where it was maintained for 2 h. After the completed dwelling time, the temperature was ramped up again at the rate of 2 °C/min until 900 °C and kept there for 1 h, after which the furnace was cooled down (5 °C/min rate) to room temperature.

An electrode-cable connection was made for the handle using conductive silver epoxy (EP-H20E, Epotek, USA) and cured according to the supplier instructions. Silver epoxy is the most practical and way to connect cables, however, it is toxic to cells. Therefore, a biocompatible and flexible styrene-ethylene-butylene-styrene (SEBS, Kraton G1650, Kraton, Belgium) elastomer coating with excellent sealing properties was applied to protect cells from silver epoxy [17]. The SEBS elastomers was dissolved in Toluene (VWR, Germany) 40% w/v and left to mix overnight. When the SEBS was completely dissolved, it was filled into a syringe and stored at room temperature and protected from light. The sealed silver epoxy and cable parts were then coated with the SEBS solution and left for curing in an oven overnight at 70 °C (Figure S1a and b in Supplementary information (SI)).

### 2.3 Structure and material properties of SU-8 and pyrolytic carbon scaffolds

The surface topography and porosity of the 3D scaffolds was characterized using scanning electron microscopy (SEM) in low-vacuum mode using a Zeiss Supra VP 40 (Carl Zeiss NTS GmbH, Germany) [11]. Image J software (National Institute of Health, Bethesda, USA) was used to determine the dimensions in the SEM images. The surface composition of the 3D printed SU-8 structures, and the 3D carbon scaffolds was characterized using a K-Alpha X-ray Photoelectron Spectrometer (XPS) system (Thermo Fisher Scientific Inc., USA) equipped with a monochromatic Al Kα X-ray source (pass energy: 200 eV for survey spectra). Data analysis was done using the Avantage software package (Thermo Fisher Scientific Inc., USA). To analyse the carbon composition of the 3D carbon scaffolds, visible Raman spectroscopy was performed using a DXR Raman Microscope (Thermo Fisher Scientific Inc., USA) at an excitation wavelength of 532 nm (10× objective and 10 mW laser power). Data processing was done using OMNIC software (Nicolet Instruments, USA). Energy Dispersive X-ray Spectroscopy (EDS) elemental distribution analysis was performed using a FEI Quanta 200 FEG-SEM coupled with an energy dispersive spectrometer (Oxford X-Max). The material nature of the carbon microflowers was further analysed using Transmission Electron Microscopy (TEM) to elucidate their morphology and crystal structure. The samples were crushed in a mortar and dispersed in ethanol on a lacey carbon-coated copper TEM grid and analyzed at 300 kV using a FEI Titan 80-300 equipped with a Gatan OneView CMOS camera and a continuum energy-loss filter.

### 2.4 Electrochemical characterization of 3D carbon scaffold electrodes

Dopamine hydrochloride (2-(3,4-dihydroxyphenyl)ethylamine hydrochloride) and cell culture tested phosphate buffered saline (PBS, pH 7.4) were purchased from Merck (Life Science, Denmark). The scaffold electrodes with cable connections were fitted into a 12-well plate (well diameter 22.1 mm depth 19 mm; NuncTM, Thermo Fisher, USA) ready to be connected to a potentiostat (Figure S1c). The electrochemical measurements were performed using a CHI 1010 potentiostat (CH Instruments, Austin TX) with a three□electrode setup. The 3D carbon scaffolds were employed as the working electrode, a platinum wire (500 µm in diameter) spiral (Advent Research materials Ltd., Oxford, England) as a counter electrode, and a Ag|AgCl (3 M KCl) (DRIREF□L, WPI, Sarasoata, FL) as the reference electrode. The 3D carbon scaffold electrodes were characterized by acquiring cyclic voltammograms (CV) in dopamine solutions (concentrations ranging between 25 nM and 250 µM) prepared in PBS, degassed by purging with nitrogen gas for 20 min. The dopamine solutions were used immediately after preparation to prevent uncontrolled oxidation of dopamine. Electrochemical characterization of the electrode scaffolds was performed after oxygen plasma treatment for 75 s using a Zepto Plasma Cleaner (Diener Electronic GmbH + Co. KG, Germany) at a pressure of 0.6 mbar and 100% power setting corresponding to 30 W. To prevent contamination of the chamber, the cables were covered with Parafilm.

### 2.5 Stem cell culture and differentiation

The 3D carbon scaffolds were prepared for cell culture by first treating them with oxygen plasma (details in section 2.4) for sterilization and to improve coating of adhesion factors and dopamine detection [15,18]. Human ventral mesencephalic neural stem cells (hVM1-Bcl-X_L_)[19,20] between passage numbers 68 and 73 were used in this study. These human neural stem cells (hNSC) were cultured for two passages after thawing from a cryovial. For experiments, the cells were seeded on poly-L-lysine (PLL, ThermoFisher Scientific) coated (10 µg/mL in PBS) 3D carbon scaffolds at a density of 10^5^ cells/scaffold. A basal medium containing DMEM/F12 with GlutaMAX (ThermoFisher Scientific), 6 g/L glucose (Sigma□Aldrich), 1% (w/v) AlbuMAX (ThermoFisher Scientific), and 5 mM HEPES (ThermoFisher Scientific) was prepared. Cells were cultured in the growth medium, which was composed of the basal medium supplemented by 1× non-essential amino acids (NEAA, Merck), 1×N□2 supplement (ThermoFisher Scientific), 1× penicillin/streptomycin mixture, as well as 20 ng/mL each of epidermal growth factor (EGF, R&D systems) and basic fibroblast growth factor (FGF, R&D systems). After 24 h incubation, the growth medium was replaced by the differentiation medium, containing the basal medium supplemented by 1× NEAA, 1× N□2 supplement, 1× penicillin/streptomycin mixture, 1 mM di-butyryl adenosine 3′, 5′□cyclic monophosphate sodium salt (dbcAMP, Merck), and 2 ng/mL human recombinant glial cell-derived neurotrophic factor (GDNF, PeproTech). After 48 h, the differentiation medium was entirely replaced by fresh medium. 2/3 of the differentiation medium was thereafter replaced with fresh differentiation medium every second day. Cells were proliferated and differentiated in a humidified incubator (HeraCell, ThermoFisher Scientific) at 37 °C in an atmosphere of 95% air/5% CO_2_. Differentiation was concluded 14 days after the first addition of the differentiation medium.

### 2.6 Characterization of hNSCs cultured and differentiated on 3D carbon scaffolds

#### Immunocytochemistry

The hNSCs, cultured and differentiated on 3D carbon scaffolds, underwent rinsing in PBS for 5 minutes, and were then fixed with paraformaldehyde (PFA) in PBS. The fixation process involved two steps: 2 % (v/v) PFA for 2 min followed by 4 % (v/v) PFA for 10 min. After thorough rinsing with PBS twice for 5 min, the cells were incubated in PBS containing 1 % (v/v) goat serum (Gibco), 1 % (v/v) horse serum (Gibco), and 0.3 % (v/v) Triton X□100 (Merck) for 1 h on a plate shaker (150 rpm) to permeabilize the cell membranes and block unspecific binding of antibodies. The cells were then incubated overnight at 4 °C with primary antibodies: rabbit polyclonal antibody against tyrosine hydroxylase (TH; 1:1000; #P40101, PelFreez Biologicals) and mouse monoclonal antibody against β-tubulin (TUBB3; 1:1000; #T8660, Merck). After completed incubation with the primary antibodies, the cells were rinsed trice for 10 min, first in PBS containing 1 % goat serum, 1 % horse serum, and 0.3 % Triton X□100, followed by by PBS containing 0.3 % Triton X□100, and finalized by PBS. After the rinsing, the cells were incubated with fluorescently labelled secondary antibodies for 30 min at room temperature: anti-mouse Alexa 546 (1:500; #A□11030, ThermoFisher) and anti-rabbit Alexa 647 (1:500; #A□21245, ThermoFisher), both generated in goat. The cell nuclei were counterstained with 2 µg/mL DAPI (1:1000; #D1306, ThermoFisher Scientific) for 10 min, followed by thorough rinsing in PBS for 10 min. Finally, the cells were mounted with an antifading solution containing 2.4 g Mowiol 4-88 (#31381, Sigma-Aldrich), 6 g glycerol (#G5516, Sigma-Aldrich), 2.5 % 1,4-Diazabicyclo(2.2.2)octane (DAPCO, #D27802, Sigma-Aldrich) in Milli-Q water and kept at 4 °C until fluorescence microscopic imaging. An inverted AxioObserver Z1 (ZeissTM, Germany) epifluorescence microscope, equipped with 5x, 10x (EC Plan-Neofluar 10x/0.30 Ph1, Zeiss™, Germany), and 20x (LD Plan-Neofluar 20x/0.4 Corr Ph2 M27, Zeiss™, Germany) objectives, was used for imaging. Images were processed using the Fiji image analysis software from NIH[21] and the Zen software package from Zeiss.

#### Scanning electron microscopy (SEM)

Cells on the 3D carbon scaffolds were rinsed in PBS for 5 min followed by fixation with 2 % (v/v) glutaraldehyde in PBS for 1 h. After thorough rinsing with PBS twice for 15 min, the cells were finally rinsed in distilled water (dH_2_O) for 15 min. The samples were then dehydrated in a series of increasing ethanol concentrations (35%, 60%, 80%, 90%, 95%, and 100%) in time intervals of 10 min each. SEM images were taken using a Carl Zeiss Auriga microscope.

### 2.7 Detection of dopamine exocytosis from cells on 3D carbon scaffolds

Prior to chronoamperometric detection of dopamine exocytosis at 600 mV (vs. Ag/AgCl (3 M KCl) RE), the cell culture medium was replaced with a baseline buffer (pH 7.4) containing 10 mM HEPES, 5 mM glucose, 1.2 mM magnesium chloride, 2 mM calcium chloride, 150 mM sodium chloride, and 5 mM potassium chloride. For measurements, a carbon scaffold with the differentiated hNSCs was dipped into a beaker containing 9 mL of baseline buffer. After recording a stable baseline current, depolarization of the cells was induced by adding 1 mL of a high-K^+^ buffer into the baseline buffer under stirring to elevate the final K^+^ concentration to 150 mM.

## 3. Results

### 3.1 Rheological characterization of fumed silica-mineral oil mixtures

Rheological measurements were used to investigate the shear-thinning behaviour of the three fused silica-mineral oil mixtures (6 to 8 % (w/v)), a critical parameter determining their suitability as a support bath for embedded 3D printing. The dynamic frequency sweep test revealed a non-Newtonian behavior of all mixtures with increased frequency. As indicated by the storage modulus (G’) crossing the loss modulus (G’’) (Figure 1a), the increased frequency led to a shift from the fluidic phase to the solid phase for all three mixtures. The dynamic strain sweep test showed that the 6 % mixture had the lowest yield strain, i.e. the point at which G’’ crosses G’, compared to the other analyzed samples (Figure 1b), which means that its viscosity and holding capacity as a support bath for embedded 3D printing was lower than for the other two mixtures. Shear rate sweeps at 10 Hz were performed by varying the shear rate from 0 to 50 Hz to determine the complex viscosity (the frequency-dependent viscosity) of the samples. The viscosity decreased as the shear rate increased for all mixtures, which means that they all behaved as a shear-thinning support (Figure 1c). Their shear-thinning behavior was also demonstrated by the transient step shear rate test. The cyclic change of strain, which is typically applied to characterize support baths for embedded printing, revealed their self-healing capacity by showing a sudden drop in viscosity at high shear, followed by a quick increase in viscosity when the applied shear was removed (Figure 1d). The temperature sweep showed that all mixtures were thermally stable until 60 °C, which is high enough for the postprocessing of SU-8 residing within the support bath (Figure 1e). Absorption spectra (Figure 1f) showed no significant difference between the mixtures in the wavelength range from 300 to 400 nm, which is the UV region needed for curing SU-8. Hence, all mixtures should be suitable for UV curing of SU-8. However, as indicated in Figure 1b, the printing trials revealed that the 6% w/v mixture had poor holding capacity, leading to the collapse of the printed structures. Both the 7 and 8 % w/v mixture showed great holding capacity until curing and release of the printed scaffolds. We proceeded with the 7 % w/v mixture as support bath for further 3D printing experiments.

**Figure 1.**
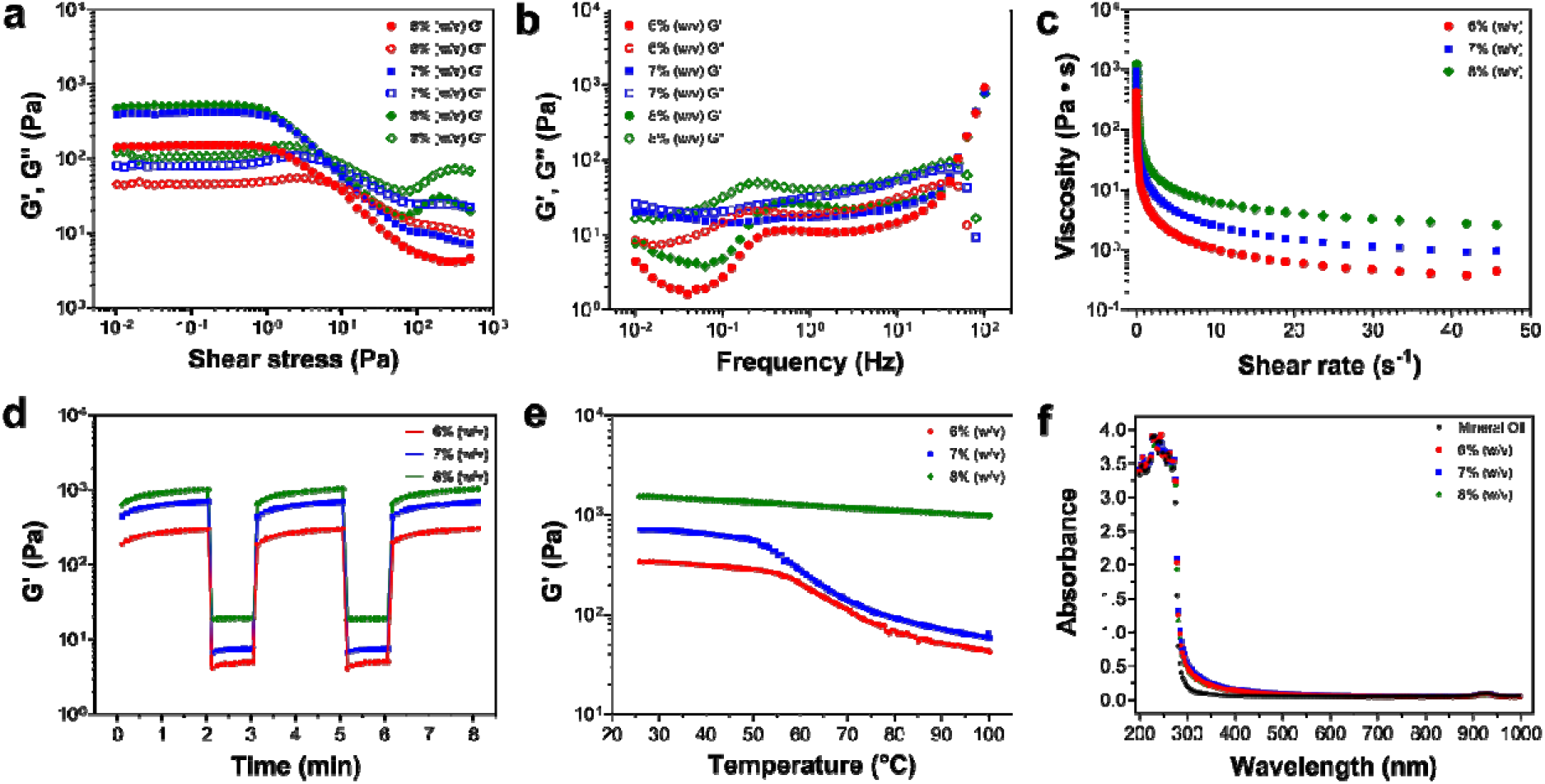
Rheological studies of three different fumed silica-mineral oil mixtures (6, 7 and 8 w/v of fumed silica in mineral oil): **a)** Strain sweep. **(b)** Frequency sweep at low 0.2 % strain. **(c)** Shear rate sweep at 10 Hz. **(d)** Cyclic strain time sweep, starting at 0.2 % strain for 120 s, then increased to 500 % strain for 60 s, and then back again to 0.2% for a new cycle. **e)** Temperature sweep conducted through the experiment. **f)** UV/Vis light absorption for the three different fused silica-mineral oil mixtures. G’: storage (or elastic) modulus and G’’: loss (or plastic) modulus.

### 3.2 Fabrication of 3D carbon scaffolds

In our previous study, we used fused filament fabrication to create polycaprolactone-based scaffolds that eliminated the formation of a necrotic core in cultured disc-shaped brain organoids [14]. In the present work, a disc-shaped scaffold structure (diameter: 15 mm) with an hexagonal honeycomb shaped infill (diameter: 2 mm, height: 1 mm, wall thickness: 0.25 mm) was designed in accordance with the previous study to maximize the scaffold surface area for cell adhesion (Figure 2a). To allow attachment of a wire for electrical connection (after pyrolysis of the SU-8 scaffold), the design was equipped with a ‘handle’ (length: 15 mm, width: 2mm) connected to the solid periphery of the scaffold structure (see section 2.2 and Figure S1a-b for details). After embedded printing of SU-8 in the 7 % (v/w) fused silica-mineral support bath, the printed structure was cured and then released from the support bath by multiple washing steps (see details in Section 2.2). After pyrolysis, the dimensions of the 3D printed SU-8 scaffolds shrank ∼50 % as shown in Figure 2b. Upon shrinkage, the 3D carbon scaffolds with the ‘handle’ for wire connection could fit in a 12-well microtiter plate (Figure S1c in the SI).

**Figure 2.**
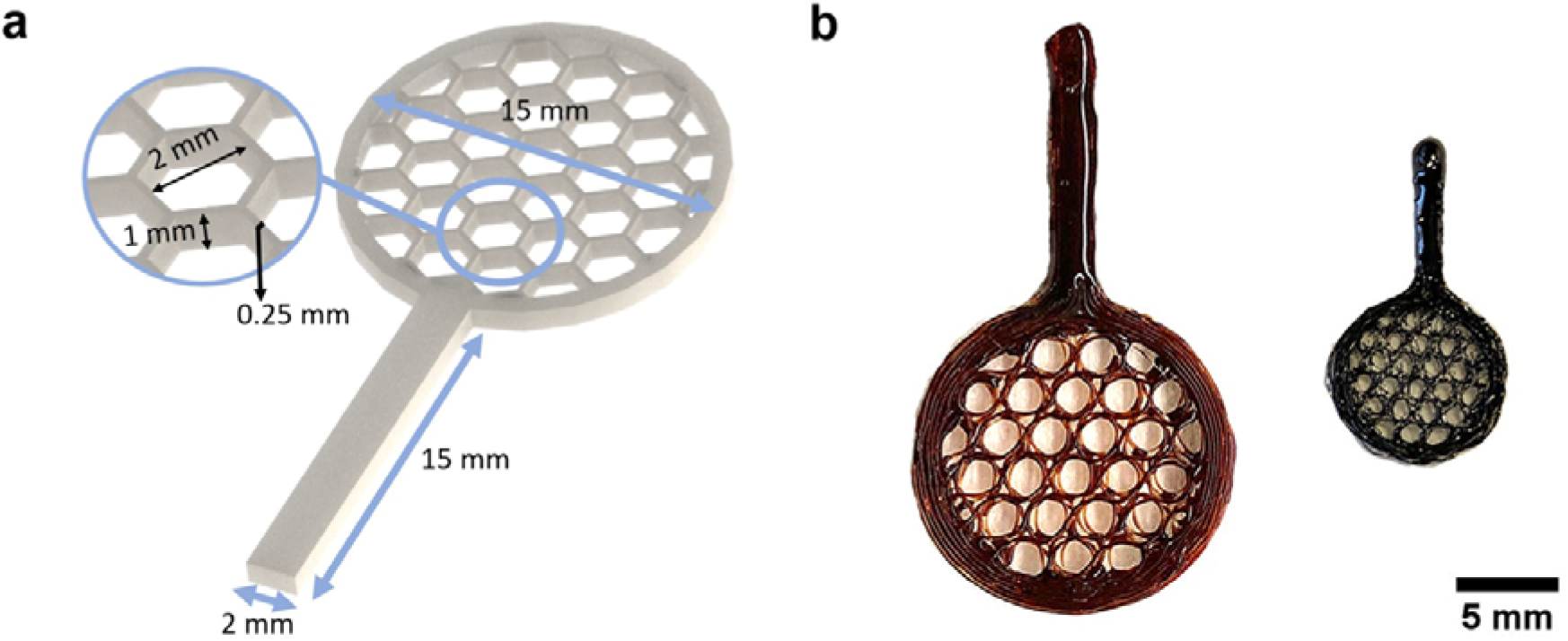
a) CAD design and dimensions of a disc-shaped scaffold consisting of an open hexagonal honeycomb lattice, and b) the embedded printed SU-8 scaffold before (left) and after pyrolysis (right), resulting in approximately 50% shrinkage of the structure after pyrolysis.

### 3.3 Morphological and physicochemical characterization of the pyrolytic carbon scaffolds

SEM was used to investigate the surface structure of the SU-8 scaffolds after embedded printing and how this structure was reproduced after pyrolysis (Figure 3). An uneven filament thickness was observed in the embedded printed SU-8 scaffolds (Figure 3a), which was reproduced in the pyrolytic carbon scaffolds (Figure 3b). This is attributed to irregular extrusion rates during printing and thus deviation from the original design (Figure 2a), which results in the formation of gaps between filament layers. The surface of the carbon structure is moreover covered by small deposits (Figure 3b), which appear as small microflowers in the magnified images (Figure 3c and d). However, we observed that the abovementioned gaps facilitated the degassing process of the scaffold segments during the pyrolytic process. Had the printed filament layers remained densely packed, the overall structure of the scaffolds might have suffered from the elevated internal pressure caused by trapped gases during pyrolysis. Gaps formed between the non-uniform filament segments increased the release rate of the gaseous byproducts during the pyrolysis process [11]

**Figure 3.**
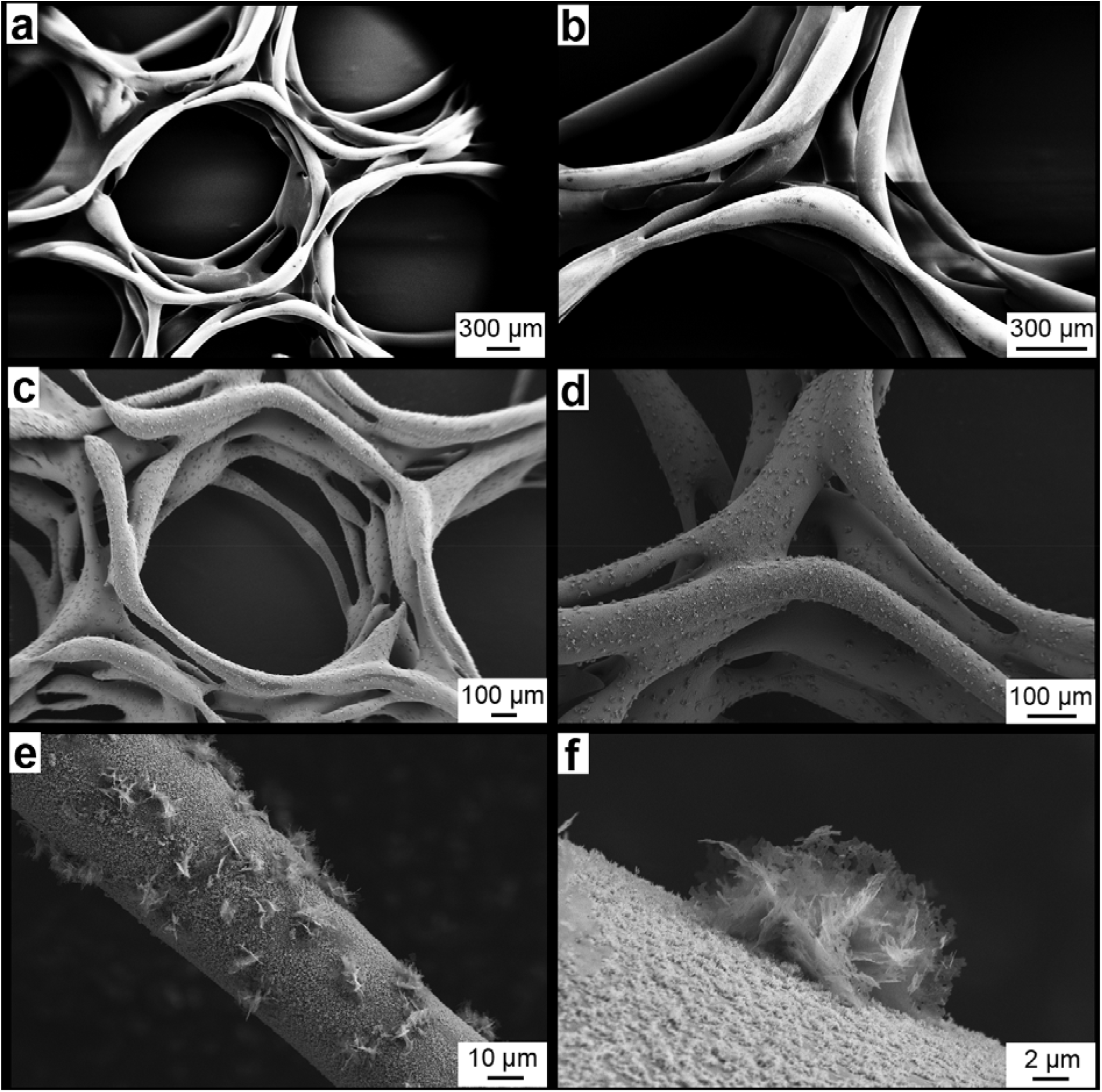
SEM images of SU-8 derived pyrolytic carbon scaffold surface: **a** and **b**) 3D printed SU-8 scaffold before pyrolysis and **c** and **d**) the corresponding structure after pyrolysis. **e**) Magnification of a pyrolyzed fiber segment showing a rough carbon surface with carbon microflowers, and **f**) magnifications of one carbon microflower.

To further characterize the surface composition of the pyrolytic carbon and especially the microflower structures, Raman, XPS, and EDS spectroscopic analyses (Figure 4), as well as TEM analysis were performed (Figure 5). Raman spectra are often used to analyze carbon materials by examining the characteristic D and G bands, which are prominent in amorphous carbons like glassy carbon. The relative intensity and shape of these bands provide valuable insights into various material properties [22]. Carbon materials with sp^2^-hybridized electrons universally exhibit the G band (E2g vibrational mode at 1580 cm□^1^). In contrast, structurally defective carbon materials, except for pure graphene, show the D band (A1g vibrational mode at 1360 cm□^1^) [11]. The presence of the D and G bands (Figure 4a) confirms that the pyrolyzed SU-8 structures have the composition characteristic of pyrolytic carbon [22]. The composition of the microflower structures were investigated by EDS, confirming that the main element was carbon (C) with small amounts of oxygen (O) and silicon (Si) also present (Figure 4b). Figure 4c presents XPS spectra of both a 3D printed SU-8 structure and the resulting carbon structure after pyrolysis. Analysis of the two spectra shows the presence of peaks around 105, 150, 390, and 530 eV, which are characteristic binding energies for silica (Si 2p), (Si 2s), carbon (C 1s), and oxygen (O 1s), respectively. The atomic ratio O/C was 88 before pyrolysis (characteristic of SU-8) and 42 after pyrolysis (see Table S1 in the SI). The presence of the Si2p peak after pyrolysis confirms the presence of remaining silica particles from the oil-silica gel bath, as also indicated by the Raman and EDS analysis. The XPS spectra reveal a reduction in the atomic ratio of silicon (Si) post-pyrolysis. This can be attributed to the volatilization and carbonization of the SU-8 matrix, which likely causes the release of any silica particles loosely adhered to the surface. Furthermore, analysis of the EDS and XPS spectra reveals a clear O peak, suggesting the presence of residual SiO_2_, likely originating from the fumed silica in the bath, which may persist even after the pyrolysis process.

**Figure 4.**
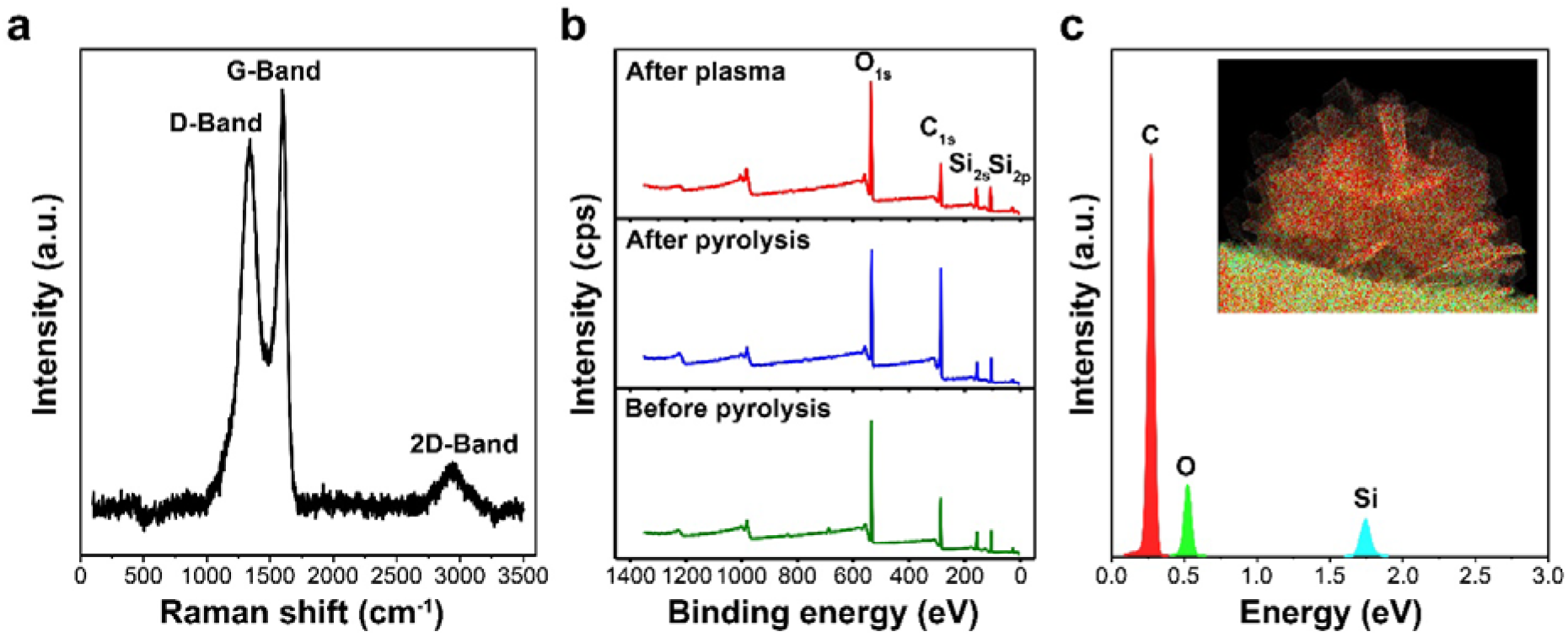
Material characterization of scaffolds: **a)** Raman spectrum for the pyrolytic carbon. **b)** Wide-range XPS spectra of SU-8 (before pyrolysis) and the SU-8 derived pyrolytic carbon (after pyrolysis and plasma). **c)** EDS spectrum (79% C, 18.3% O, 2.5% Si) for elemental mapping of carbon microflower structures (inset).

**Figure 5.**
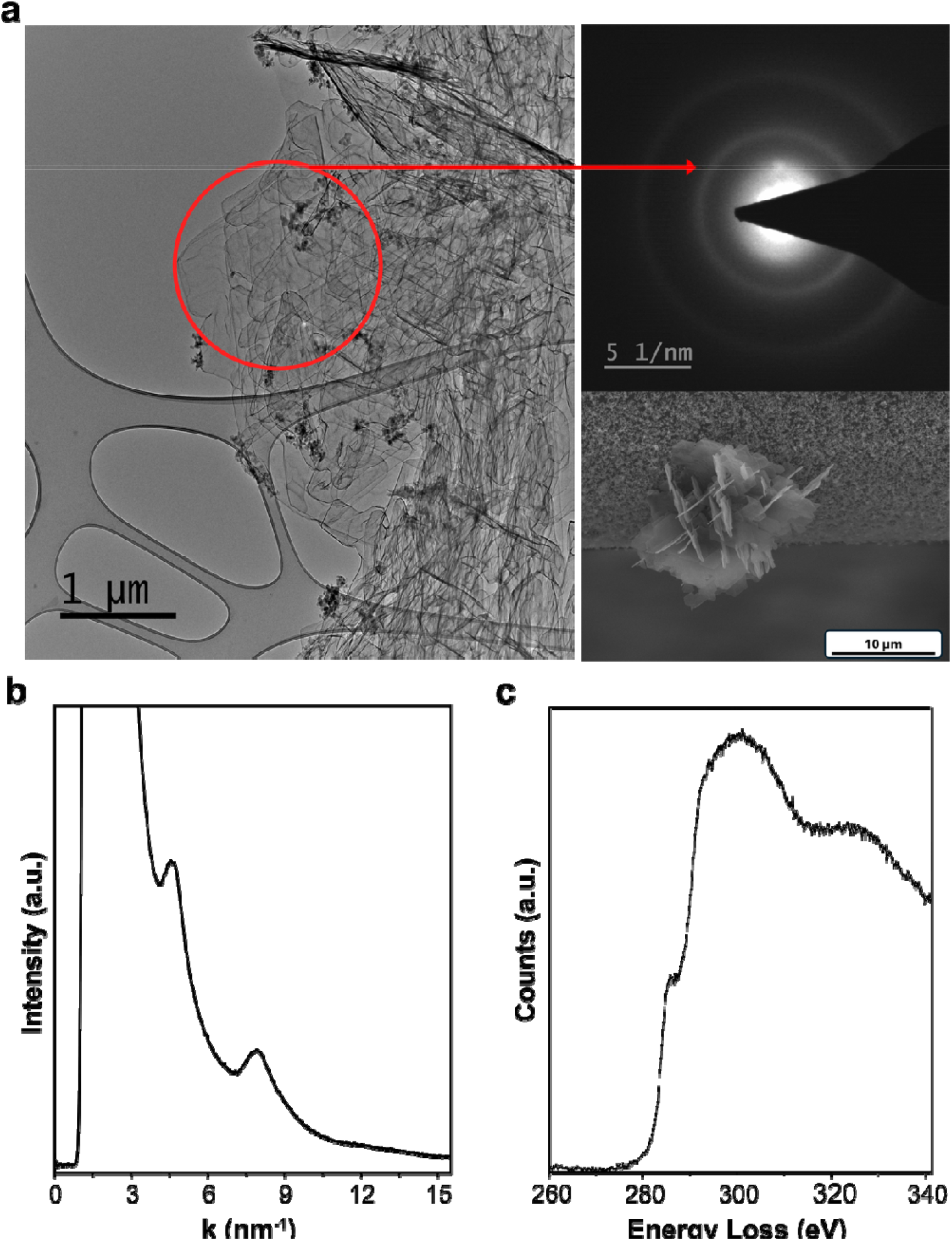
**a)** TEM image (*left*) and diffraction pattern (*top right*) of the thin carbon leaves of the microflower shown in the secondary SEM image (*bottom right*) acquired at 3 kV. **b)** Rotational average of an electron diffraction pattern acquired from the red circular region indicated in a) showing peaks corresponding to graphene (010) and (110) planes. **c)** Electron energy loss spectrum showing the characteristic K-edge at 284 eV acquired from the same region as the diffraction pattern in a).

TEM of the calcined samples show plate-like structures of the carbon microflowers on the surface of the filaments (SEMs in Figure 3e and f) about 200 – 1000 nm in diameter (Figure 5a). The edge contrast indicates a thickness of these structures of a few tens of nm [23]. Electron diffraction of individual structures shows reflections at 4.4 nm^-1^ corresponding to the (010) planes at 2.1 Å and 7.8 nm^-1^ corresponding to the (110) reflection of graphite at 1.2 Å (Figure 5b), indicating a graphene-like structure [24]. The (002) reflection is absent in the patterns indicating little ordering along the C axis. Electron energy-loss spectroscopy (Figure 5c) confirms the carbon chemistry and the lack of ordering along the C axis.

### 3.4 Electrochemical characterization of 3D carbon scaffolds

The 3D carbon scaffolds were first treated with oxygen plasma and then characterized using cyclic voltammetry to evaluate their ability for oxidation of dopamine, and hence their suitability for detection of dopamine exocytosis. The CVs for 250 µM dopamine at different scan rates (Figure 6a) show that the separation between the anodic and cathodic peaks increases with the scan rate, which is typical for quasi-reversible electrochemistry. The linear relationship between the anodic peak current and the square root of the scan rate (Figure 6b) indicates that the conductivity is not a limiting factor even at higher scan rates [25]. The 3D carbon scaffolds were also assessed in terms of their sensitivity to dopamine detection by acquiring CVs at different dopamine concentrations (Figure 6c), showing that the anodic peak currents are proportional to dopamine concentration with a calculated sensitivity of 1 μA/μM.

**Figure 6.**
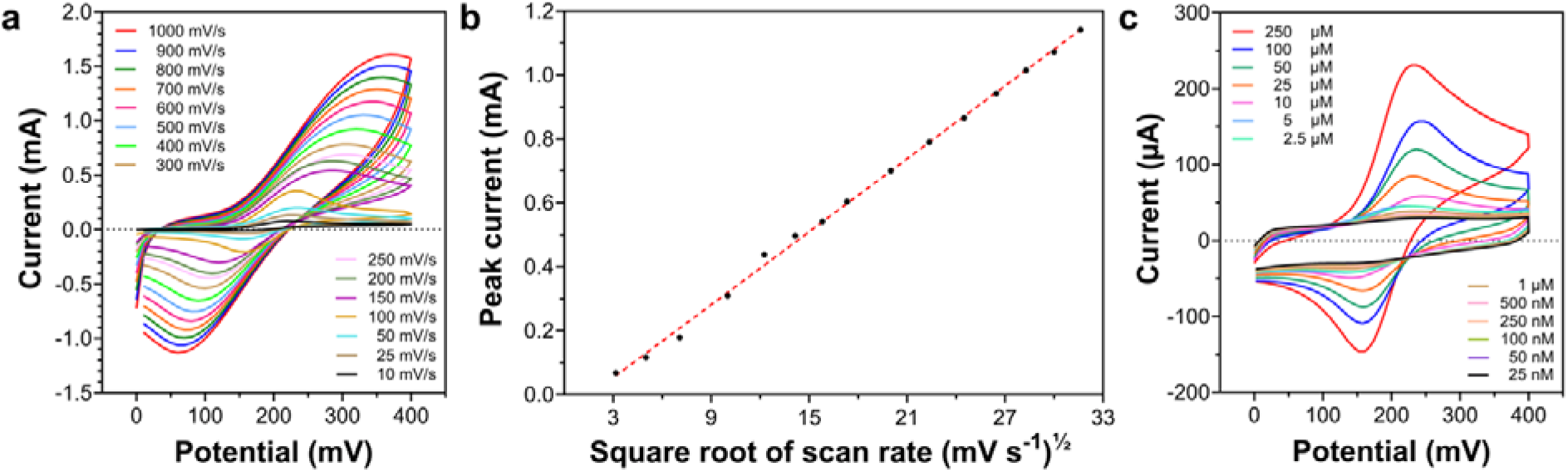
Cyclic voltammetric characterization of 3D carbon scaffolds using dopamine: **a)** CVs for 250 μM dopamine in PBS at different scan rates; **b)** Peak current vs. square root of scan rate for the CVs in a). **c)** CVs for varying concentrations of dopamine in PBS (scan rate: 100 mV/s).

To test how the 3D carbon scaffolds respond to transient injections of dopamine, we performed amperometric measurements by repeated 25 µL injections of a 250 µM dopamine solution to the center of the scaffold using a syringe pump. Typical amperometric responses from a 3D carbon scaffold clearly show that the 3D carbon scaffold reproducibly responds to transiently introduced dopamine (RSD 1.9%) (Figure S2).

### 3.6 Differentiation of hNSCs on 3D carbon scaffolds and detection of dopamine exocytosis

We have previously shown that SU-8-derived pyrolytic carbon is an efficient substrate for the differentiation of human neural stem cells (hNSCs) into dopaminergic neurons, at the same time functioning as an electrode for the detection of dopamine exocytosis [15,25,26]. The morphology and identity of the hNSCs after 14 days of differentiation on the 3D carbon scaffolds was investigated by SEM and immunocytochemistry [26]. hNSCs were cultured for 24 h at an initial density of 10^5^ cells/scaffold on oxygen plasma treated and PLL-coated carbon scaffolds. The PLL coating was applied to create a positively charged surface, promoting uniform cell adhesion. The morphology of the cells was assessed after 14 days of differentiation and showed effective differentiation with typical neuronal morphology. It can be seen that the hNSCs differentiate on the scaffold surface, on the microflowers, and across the gaps of different filaments, as evidenced by SEM images (Figure 7). Immunocytochemistry (Figure 8) confirms the neural (green, TUBB3+ cells, Figure 8a and d) and dopaminergic phenotype (red, TH+ cells, Figure 8b and e). TH is the rate-limiting enzyme of dopamine biosynthesis, and biomarker for dopamine positive cells. These results were consistent with our previous studies, i.e., the pyrolytic carbon material sustains differentiation into neuronal and dopaminergic phenotype.

**Figure 7.**
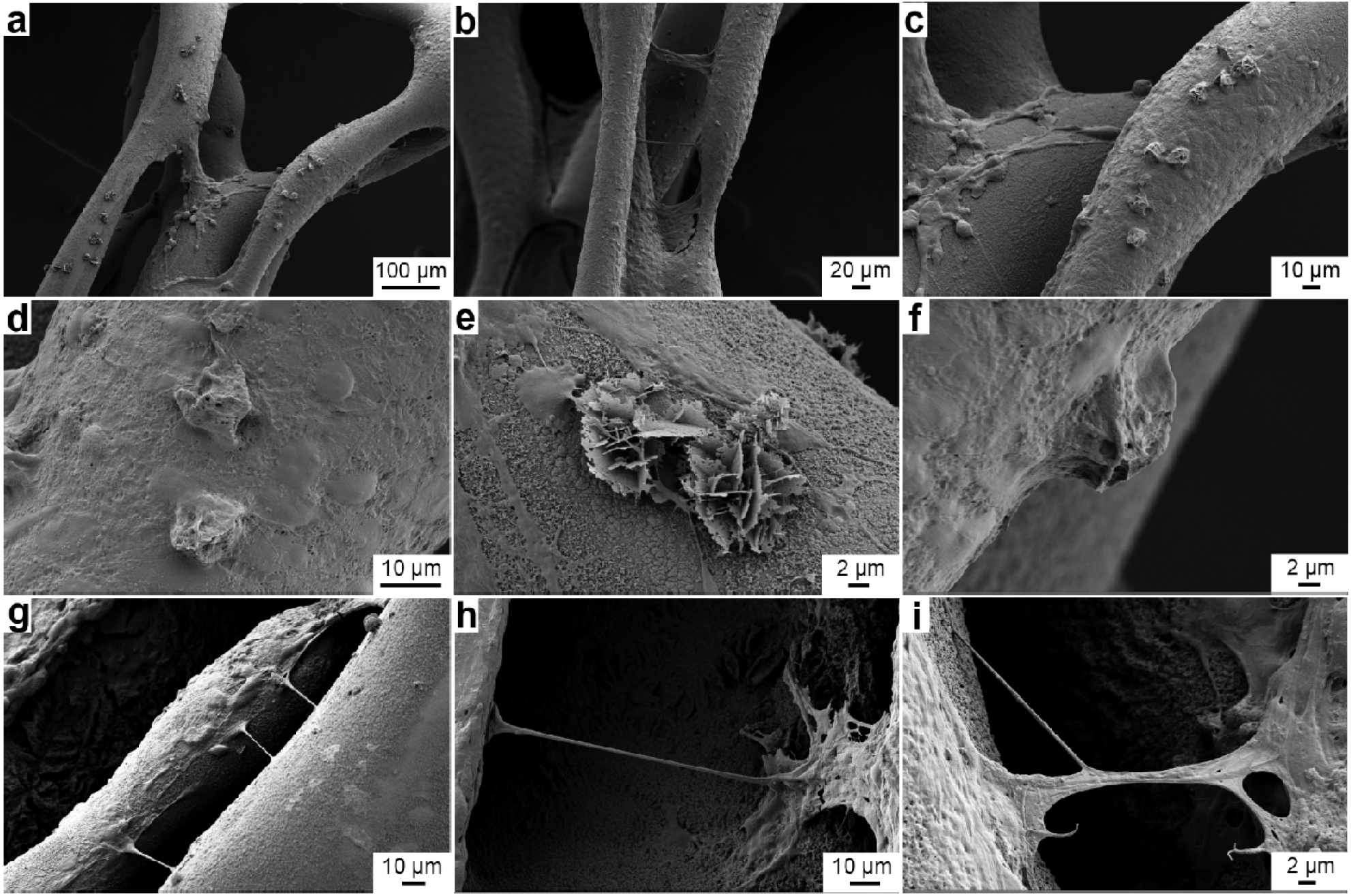
SEM images of differentiating hNSCs on a carbon scaffold electrodes at different magnifications: Morphology of hNSCs after 14 days of differentiation.

**Figure 8.**
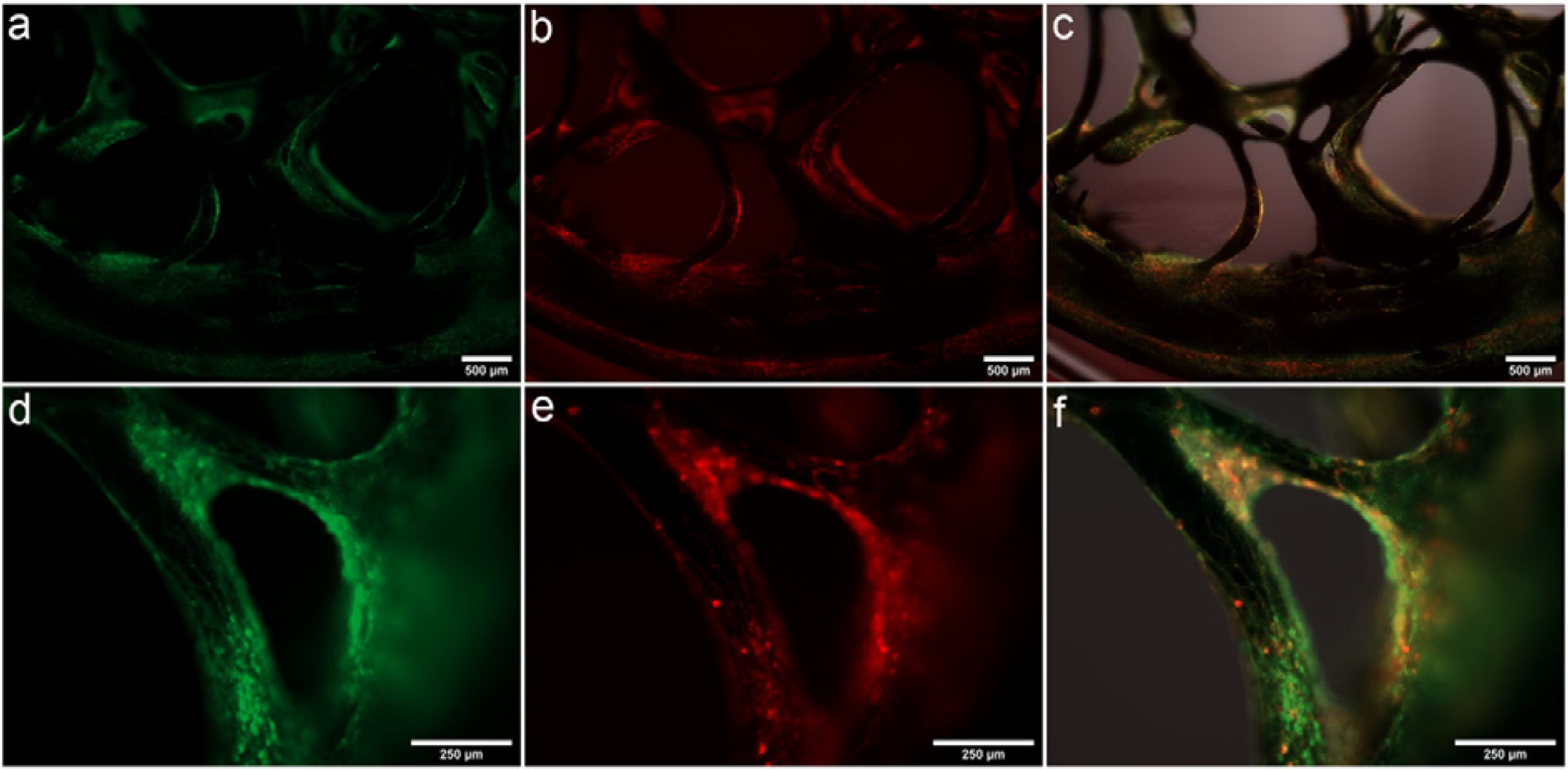
Immunocytochemistry of hNSCs differentiated on 3D carbon scaffolds at two different magnifications. **a** and **d**) β-tubulin (green). **b** and **e**) TH (red). **c** and **f**) Merge (green and red).

Detection of dopamine exocytosis from the differentiated hNSCs on the carbon scaffolds was conducted by first immersing the cell scaffolds in the baseline buffer and recording the current baseline. Depolarization to trigger dopamine exocytosis was then induced by increasing the concentration of K^+^ [15,25]. Figure 9 shows a typical amperometric current response (blue trace) recorded on a scaffold with hNSCs after 14-days of differentiation, as well as a control measurement with a scaffold containing non-differentiated cells (orange trace) (for additional exocytosis measurements, see Figure S3. The presented amperometric recordings show that the 3D carbon scaffolds were suitable for electrochemical detection of dopamine exocytosis.

**Figure 9.**
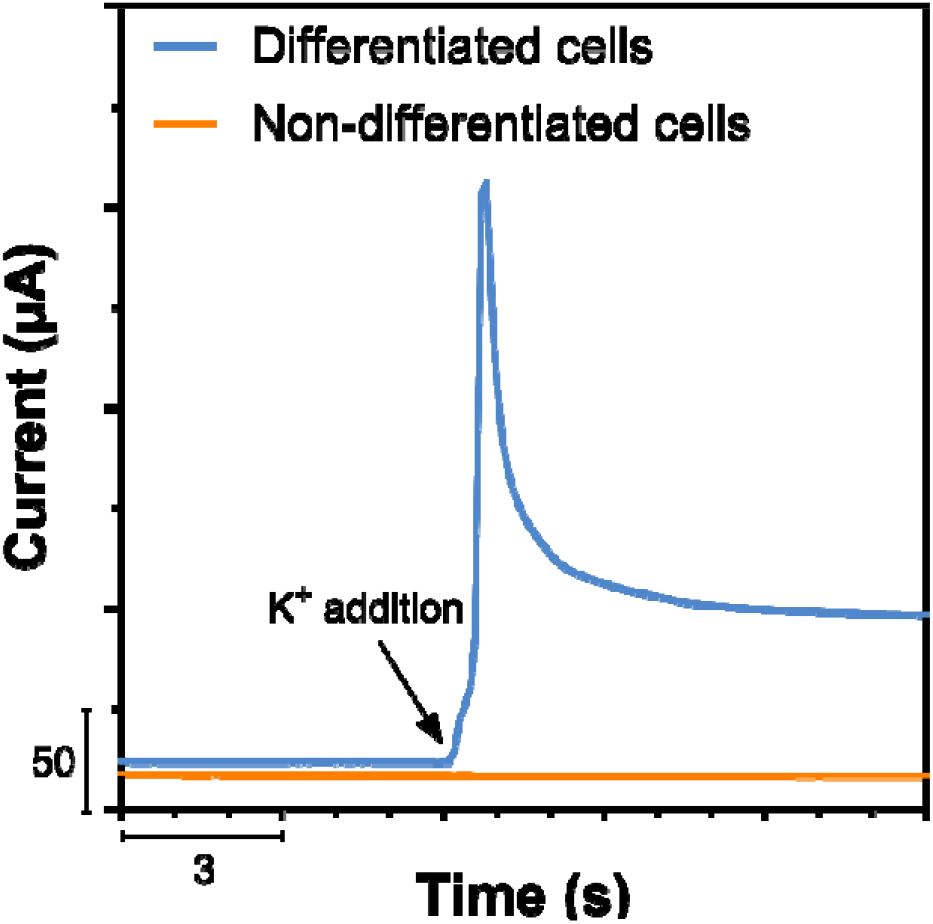
Amperometric exocytosis measurements after depolarization (150 mM K^+^) of 14-days differentiated hNSCs on 3D carbon scaffold (blue trace) as compared with a scaffold containing non-differentiated hNSCs (orange trace).

## 4. Discussion

The present study demonstrates that embedded printing of SU-8 within a fumed silica-mineral oil support bath followed by pyrolysis provides a practical route for fabrication of electrically conductive three-dimensional carbon scaffold electrodes. The rheological characterization showed that support baths containing 7–8% (w/v) fumed silica exhibited sufficient yield stress, shear-thinning behavior, and rapid viscosity recovery to support embedded printing of low-viscosity SU-8. These properties are essential for maintaining structural fidelity during printing while still allowing nozzle movement through the support medium [12,13].

A notable feature of the fabrication process was the preservation of the overall scaffold architecture following pyrolysis despite an approximately 50% dimensional shrinkage. Such shrinkage is expected during conversion of SU-8 into pyrolytic carbon due to the removal of non-carbon elements and densification of the carbon network[11] . Importantly, the shrinkage occurred without compromising the open honeycomb structure, allowing the resulting scaffold electrodes to maintain a highly porous architecture suitable for both cell culture and electrochemical sensing.

Morphological characterization revealed the presence of carbon microflowers decorating the surface of the pyrolyzed filaments. Raman spectroscopy confirmed the characteristic D and G bands of pyrolytic carbon, indicating a defect-rich carbon material containing substantial sp^2^ bonding [22]. The slightly higher G-band intensity relative to the D-band suggests a relatively high degree of graphitic character while maintaining sufficient disorder to generate electrochemically active sites. Such defect-rich carbon materials are generally advantageous for electrochemical sensing due to their ability to facilitate rapid electron transfer reactions[1].

The combined EDS, XPS and TEM analyses provided further insight into the nature of the carbon microflower structures. Residual silicon-containing species originating from the fumed silica support bath were detected after pyrolysis. It is therefore plausible that remaining silica particles acted as nucleation sites during carbonization, promoting localized growth of the microflowers. TEM diffraction patterns displaying reflections corresponding to graphene (010) and (110) planes further indicate that these structures possess graphene-like ordering, although the absence of the graphite (002) reflection suggests limited ordering along the c-axis[23,24]. Together, these observations indicate that pyrolysis generated hierarchical surface features with nanoscale graphitic characteristics.

The electrochemical studies demonstrated that the scaffold electrodes are suitable for dopamine sensing. The linear dependence of peak current on the square root of scan rate suggests diffusion-controlled electrochemical behavior, while the quasi-reversible voltammetric response is consistent with dopamine oxidation at carbon electrodes. The relatively high sensitivity obtained for dopamine detection may be attributed to the combination of the intrinsically porous scaffold architecture, rough filament surfaces, and the presence of graphene-like microflower structures, which collectively increase the electrochemically active surface area. In addition, the highly defective carbon structure evidenced by Raman spectroscopy likely contributes to efficient electron-transfer kinetics [1] .

Beyond their electrochemical functionality, the scaffold electrodes successfully supported attachment, growth and differentiation of human neural stem cells into a dopaminergic phenotype. SEM imaging demonstrated extensive cellular coverage throughout the scaffold, including on the microflower-decorated surfaces and across gaps between neighboring filaments. Immunocytochemistry confirmed expression of both the neuronal marker β-tubulin III and the dopaminergic marker tyrosine hydroxylase. These findings agree with previous studies demonstrating the favorable interactions between pyrolytic carbon surfaces and neural stem cells [15] and indicate that the embedded-printing fabrication route does not compromise the biological compatibility of the resulting carbon material.

A key outcome of this work is the integration of structural support, electrochemical sensing and neural differentiation within a single platform. Following potassium-induced depolarization, the scaffold electrodes were able to record dopamine release from differentiated cells in real time, whereas non-differentiated controls did not produce comparable responses. This demonstrates that the scaffold functions not only as a cell culture substrate but also as an active bioelectronic interface. Such dual functionality may be particularly valuable for future engineered neural tissues and brain organoid systems, where continuous monitoring of neurotransmitter release could provide insights into maturation, functionality and disease-related phenotypes.

Overall, the combination of embedded printing and pyrolysis offers a scalable strategy for generating architecturally complex carbon biointerfaces. The resulting graphene-like pyrolytic carbon scaffold electrodes provide a unique combination of mechanical support, neural compatibility and electrochemical sensing capability, making them promising candidates for future bioelectronic tissue-engineering applications.

## 5. Conclusion

In summary, we present a straightforward embedded 3D-printing approach based on a shear-thinning support bath containing 7% (w/v) fused silica nanoparticles in mineral oil for the fabrication of complex three-dimensional architectures. Following pyrolysis, the printed structures are converted into conductive carbon scaffolds with a rough surface morphology decorated by graphene-like microflowers. These scaffolds simultaneously serve as substrates for neural cell growth and differentiation and as electrochemical sensing platforms. The resulting 3D carbon scaffolds exhibited excellent electrochemical performance, enabling dopamine detection down to 25 nM. Furthermore, they supported the culture and differentiation of human neural stem cells into a dopaminergic phenotype while allowing real-time monitoring of dopamine release. Collectively, these findings establish the proposed platform as a promising tool for neuronal biointerfaces, with future potential for electrically guided neural differentiation and the generation of disc-shaped brain organoids.

## Acknowledgements

This project has received funding from the European Union’s Horizon 2020 research and innovation program under the Marie Skłodowska-Curie grant agreement No. 722779 and was conducted within the “Training 4 Cell Regenerative Medicine” (T4CRM) network. Work at DTU Health Tech was supported by grant (to A.A.D. and M.D.), “Organoids for Virus Research” (OrganoVIR) European Union Horizon 2020 Programme (H2020□MSCA□ITN□2018) under the Marie Skłodowska□Curie Innovative Training Network and Grant Agreement No. 812673. We acknowledge the OpenMIND project (HORIZON-EIC-2021-PATHFINDEROPEN-01 Project 101047177) for providing essential infrastructure and support that contributed to this researcsh.

## Ethics statement

Approval for using human neural stem cells has been obtained from the Danish authority (Arbejdstilsynet).

