## Supplementary for "Embedded Printing-Derived Pyrolytic Carbon Electrodes with Graphene-like Features for Neural Stem Cell Differentiation and Dopamine Sensing"

#### ^1^Department of Biotechnology and Biomedicine, Technical University of Denmark, Kgs. Lyngby, 2800, Denmark

#### ^2^Department of Health Technology, Technical University of Denmark, Kgs. Lyngby, 2800, Denmark

*^3^Department of Nanolab, Technical University of Denmark, Kgs. Lyngby, Denmark*

*^4^ Electronic Engineering Department, University of Rome Tor Vergata, Via del Politecnico 1, 00133 Rome, Italy*

#### ^5^Novo Nordisk Foundation Center for Stem Cell Medicine (reNEW)), Faculty of Health and Medical Sciences, University of Copenhagen, Copenhagen, Denmark

#### ^6^Department of Molecular Biology, Autonomous University of Madrid, Center of Molecular Biology Severo Ochoa (CBMSO, UAM-CSIC), Madrid, Spain

#### ^7^Center for Analysis and Synthesis, Department of Chemistry, Lund University, Lund, Sweden

*Corresponding authors:

*Giorgio Scordo:*

*Jenny Emneus*:

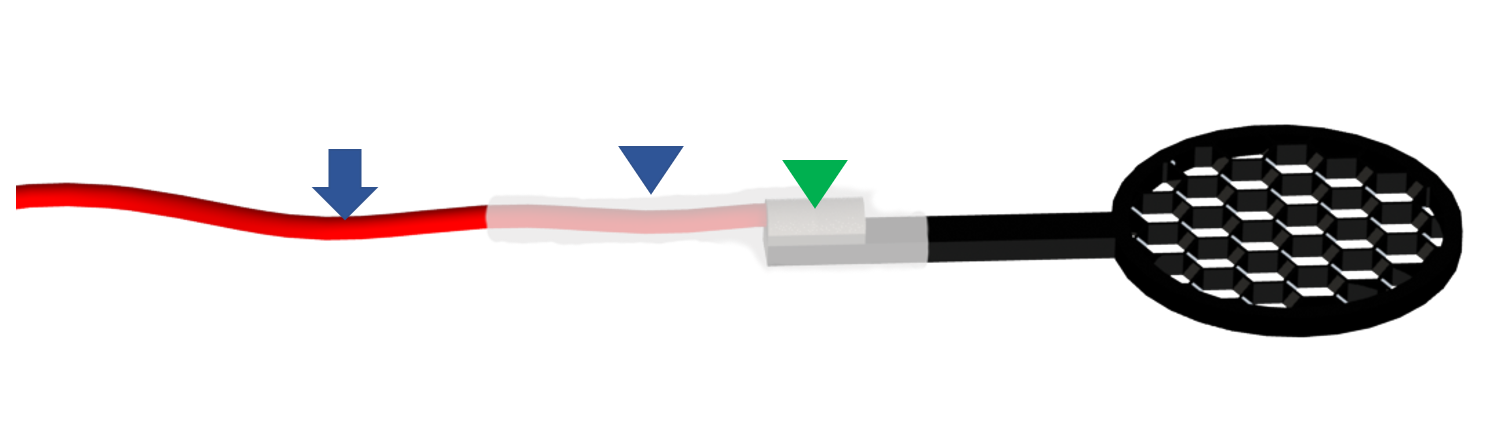

**a**

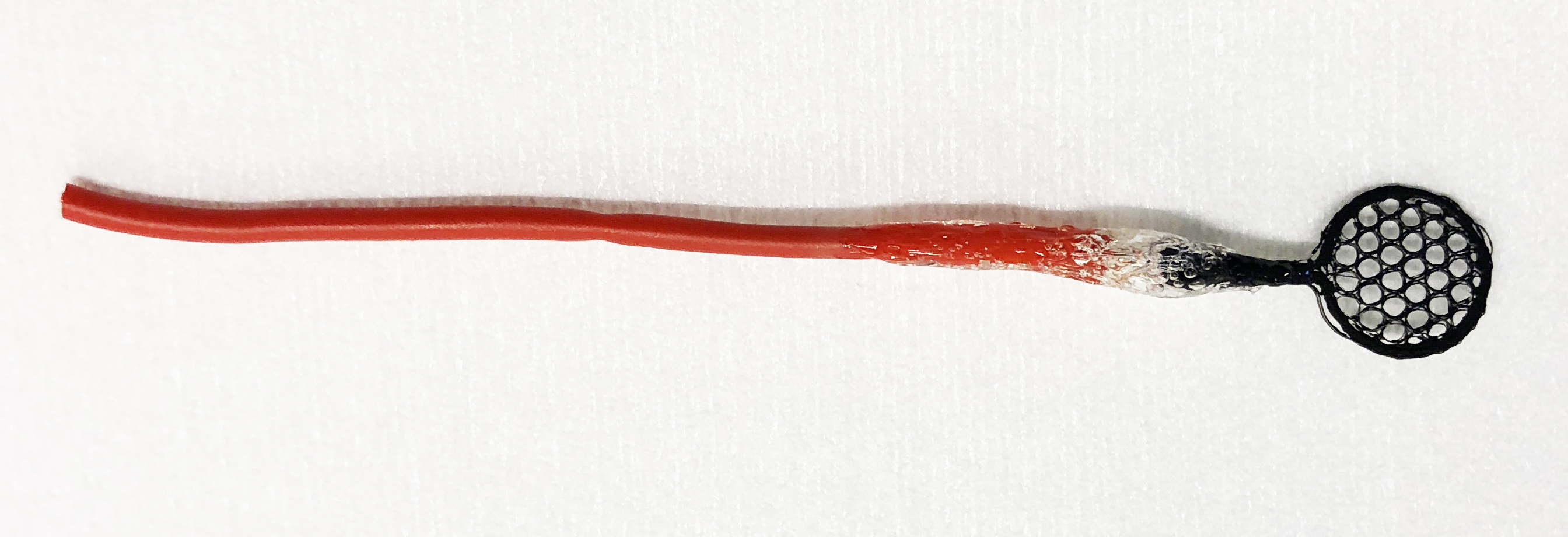

**b**

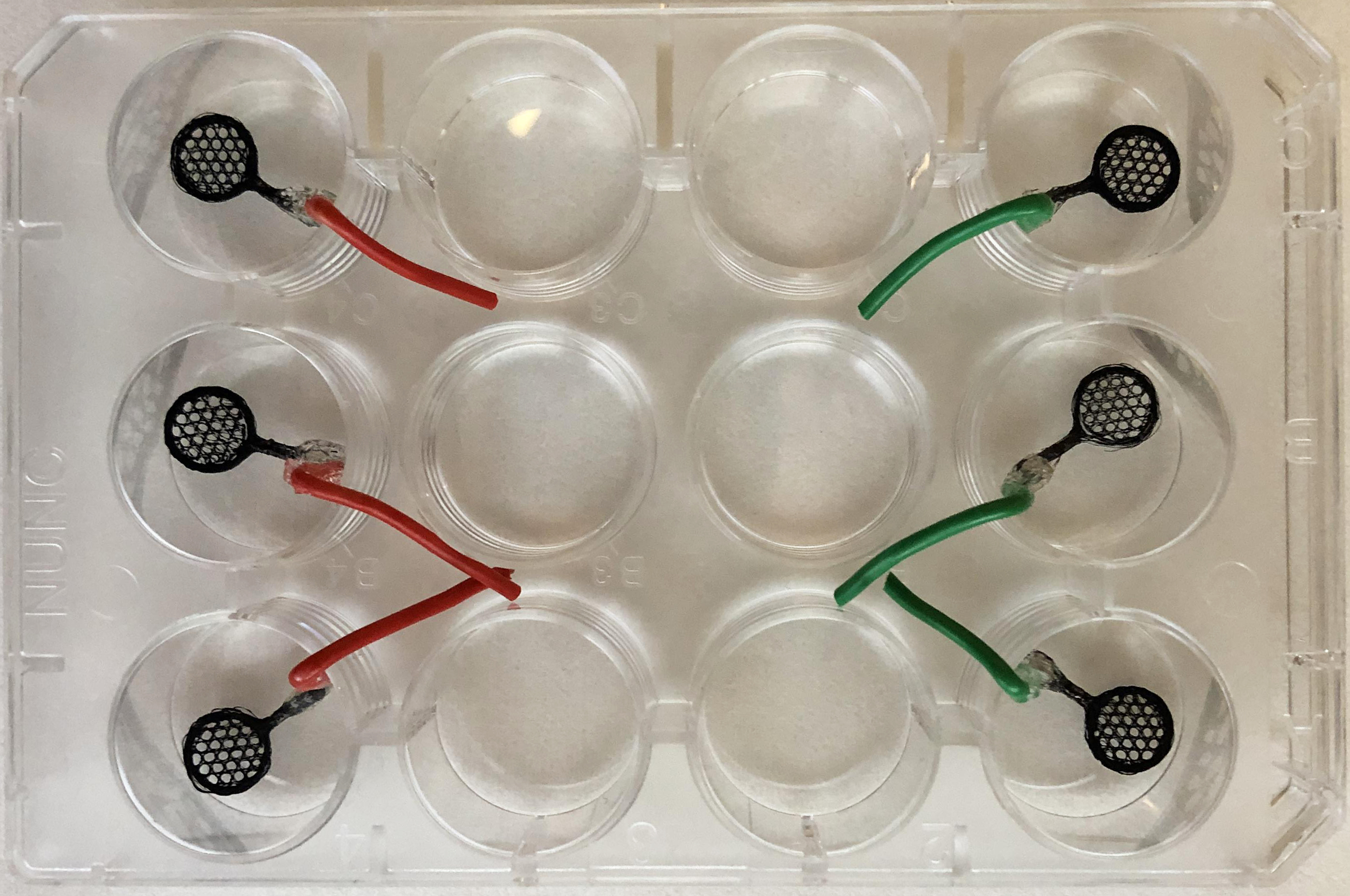

**c**

**Figure S1:** Experimental setup. **a)** Illustration of the fully prepared electrodes for detection (Blue arrow indicates Cable, Blue arrowhead indicates SEBS coating, green arrowhead indicates silver epoxy), b) electrode after curing and coating steps, c) experimental setup for cell culture.

**Table S1** Surface atomic composition measured by XPS before (SU-8) and after pyrolysis process and after O_2_ plasma treatment.

| **Material/treatment** | **Atomic composition (%)** | | | **O/C ratio (%)** | **Atomic composition (%)**  **Approximation for C & O**  (SiO_2_ contribution subtracted) | |
| --- | --- | --- | --- | --- | --- | --- |
|  | **C** | **O** | **Si** | **Based on XPS** | **C** | **O** |
| 3D printed SU-8 before pyrolysis | 42.6 | 37.5 | 17.5 | 88.0 | 94^1^ | 6^1^ |
| Pyrolytic carbon | 61.9 | 26.0 | 12.1 | 42.0 | 97^2^ | 3^2^ |
| Pyrolytic carbon after O_2_ plasma | 36.2 | 44.7 | 19.1 | 123 | 85^3^ | 15^3^ |

**Approximated atomic composition for C and O after subtraction of SiO_2_ contribution**

^1^**3D printed SU-8 before pyrolysis**

The 3D printed SU-8 is covered by silica nanoparticles. Assuming SiO_2_ composition with an O/Si ratio of 2:1, the 37.5% oxygen on the surface comprises roughly 2 x 17.5% = 35% oxygen from the silica. This means that 42,6% carbon and ~2,5% oxygen of the total composition are from SU-8, leading to an approximate composition of SU-8 as ~94% carbon and ~6% oxygen. The oxygen content of SU-8 should be around 16-19% in the bulk (Astley et al, 2025- DOI: [10.1016/j.carbon.2025.120728](https://doi.org/10.1016/j.carbon.2025.120728)), which means that the approximated oxygen content of SU-8 is clearly lower than expected based on studies relying on pristine SU-8.

Considering that the typical penetration depth of X-rays in XPS experiments is only a few nanometers, the SU-8 surface may be effectively shielded by the silica nanoparticles with sizes in the range of a few tens of nanometers. Hence, the low approximate oxygen content is a result of incomplete detection of the oxygen signal originating from SU-8.

^2^**Pyrolytic carbon**

Based on the same reasoning as presented above, the silica nanoparticles remaining on the formed pyrolytic carbon may contribute to 24.2% of the total detected oxygen content of 26.0%. This means that 61.9% carbon and 1.8% oxygen of the total composition represent the pyrolytic carbon. An approximate carbon and oxygen composition of the pyrolytic carbon

would be ~97% carbon and ~3% oxygen, which is similar to what other studies have reported (Peltola et al., 2017-DOI: [10.1039/c7tb02469j](https://doi.org/10.1039/c7tb02469j); Hemanth et al., 2017-DOI: [10.1016/j.carbon.2017.05.090](https://www.sciencedirect.com/science/article/pii/S000862231730547X?via%3Dihub); Mendez-Linan et al., 2020-DOI: [10.1016/j.carbon.2020.06.085](https://doi.org/10.1016/j.carbon.2020.06.085))

Again, after pyrolysis, the presence of silica nanoparticles on the surface can shield the underlying carbon. However, due to the diffusion of silica nanoparticles into the carbon matrix at high temperatures, along with the flow of softened organic materials toward the surface, less silicon is detected. Consequently, more carbon is observed within the few nanometers probed by XPS on the sample surface.

^3^**Pyrolytic carbon after O_2_ plasma**

Due to presence of silica nanoparticles, of the 44.7% oxygen on the surface, 38.2% can be attributed to SiO_2_. This means that 36.2% carbon and 6.5% oxygen of the total composition can be assigned to O_2_ treated pyrolytic carbon, which corresponds to the approximate composition of 85% carbon and 15% oxygen. This increase in oxygen content is explained by formation of oxygen surface groups by the plasma treatment (Peltola et al., 2017-DOI: [10.1039/c7tb02469j](https://doi.org/10.1039/c7tb02469j); Mendez-Linan et al., 2020-DOI: [10.1016/j.carbon.2020.06.085](https://doi.org/10.1016/j.carbon.2020.06.085); Pankratova et al., 2022-DOI: [10.1016/j.electacta.2022.139987](https://doi.org/10.1016/j.electacta.2022.139987)).

In XPS analysis, the outermost surface layer largely determines the detected elemental composition. While the plasma treatment does not alter the silica nanoparticles, it can introduce oxygen-containing functional groups to the carbon. Therefore, an increase in oxygen atomic percentage on the surface is expected due to carbon functionalization.

Based on the XPS results, two general conclusions can be drawn:

1. Presence of silica nanoparticles on the surface after 3D printing: These may have catalyzed the pyrolysis process, contributing to the formation of carbon micro-flowers.
2. Effect of plasma treatment: The increase in oxygen content indicates that carbon has been functionalized by the plasma treatment. This suggests that carbon is accessible at the surface, i.e., the surface is not entirely covered by silica nanoparticles. Instead, we observe a composite surface consisting of both carbon and silica.

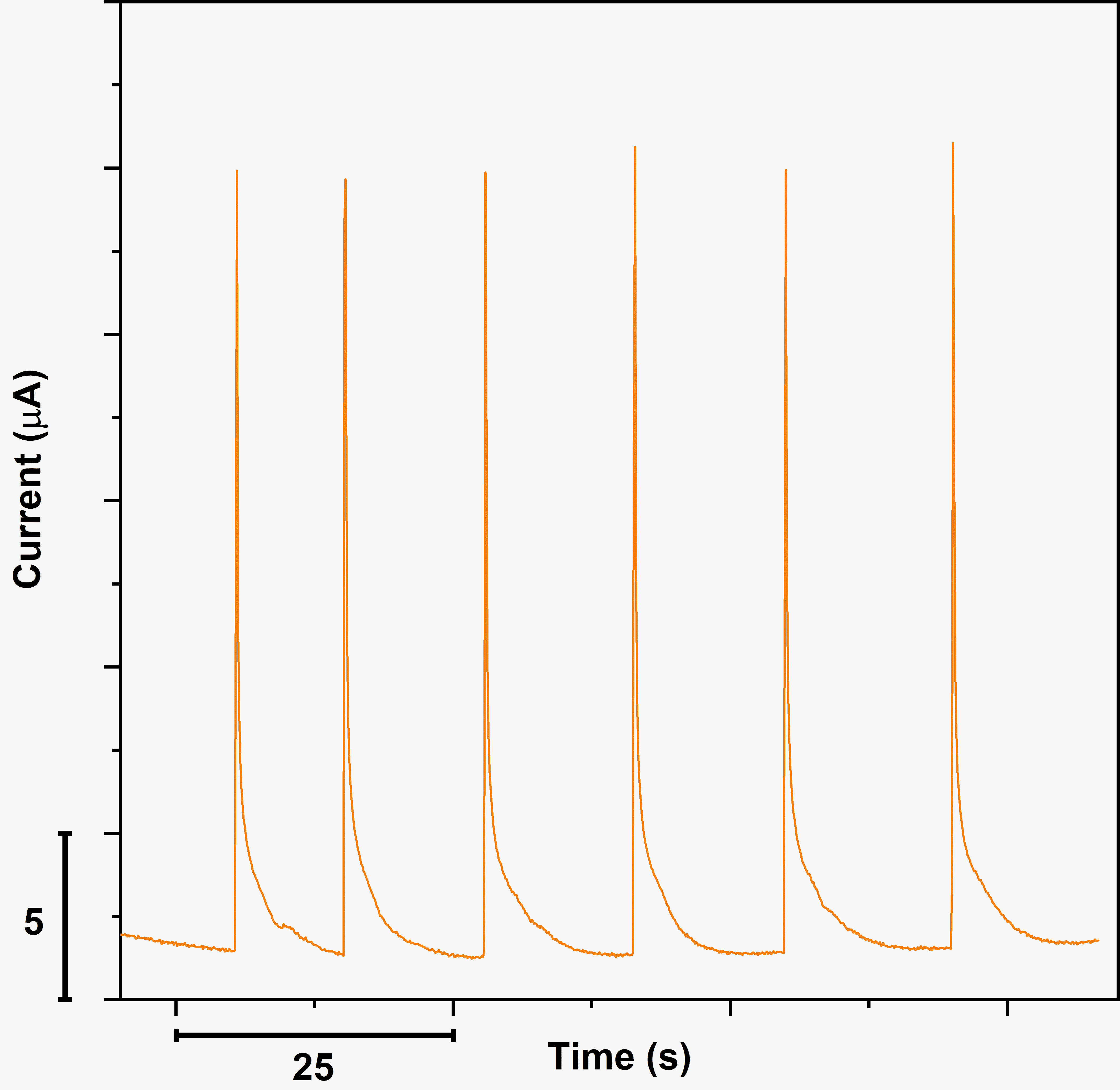

**Figure S2.** Chronoamperometry: Six repeated 25 µL injections of 250 µM dopamine.

| 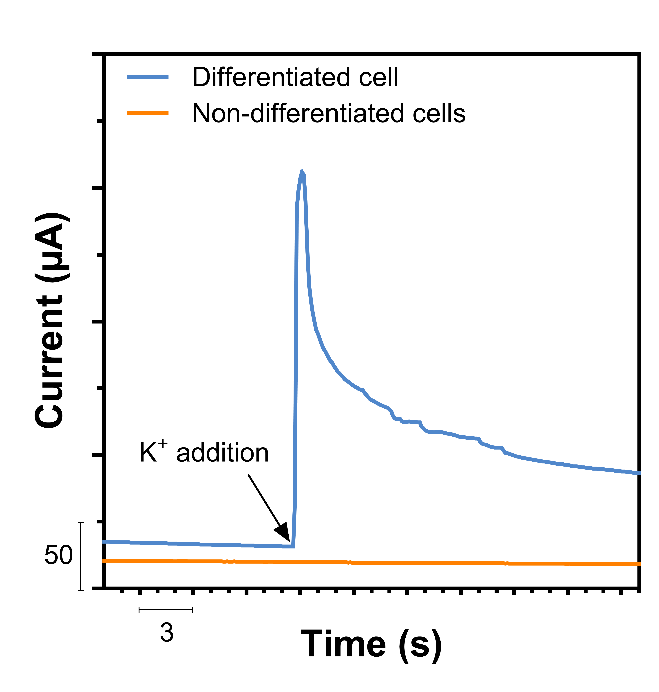 | 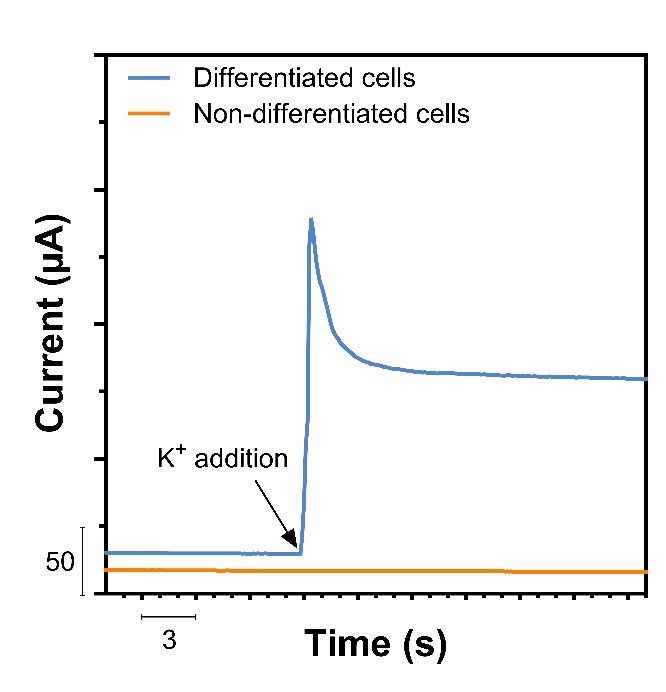 |
| --- | --- |

**Figure S3.** Amperometric exocytosis measurements recorded from two additional carbon scaffolds (compare Figure 9) revealing distinct differences between differentiated (blue lines) and non-differentiated (orange lines) human neural stem cells (hNSCs) when stimulated with K^+^ ions (final concentration ∼150 mM).
